# Single-cell meta-analysis of T cells reveals clonal dynamics of response to checkpoint immunotherapy

**DOI:** 10.1101/2024.09.25.614877

**Authors:** Ofir Shorer, Asaf Pinhasi, Keren Yizhak

## Abstract

Despite the crucial role of T cell clones in the anti-tumor activity, their characterization and association with clinical outcome following immune checkpoint inhibitors (ICI) is lacking. Here we analyzed paired single-cell RNA-sequencing/T-cell receptor sequencing of 767,606 T cells from 460 samples spanning 6 cancer types. We found a robust signature of response based on expanded CD8^+^ clones that differentiates between responders and non-responders. Analysis of persistent clones showed transcriptional changes that are differentially induced by therapy in the different response groups, suggesting an improved reinvigoration capacity in responding patients. Moreover, a gene trajectory analysis revealed changes in the pseudo-temporal state of de-novo clones that are associated with response to therapy. Lastly, we found that clones shared between tumor and blood are more abundant in non-responders and execute distinct transcriptional programs. Overall, our results highlight differences in clonal transcriptional states that are linked to patient response, offering valuable insights into the mechanisms driving effective anti-tumor immunity.

**Graphical abstract:** 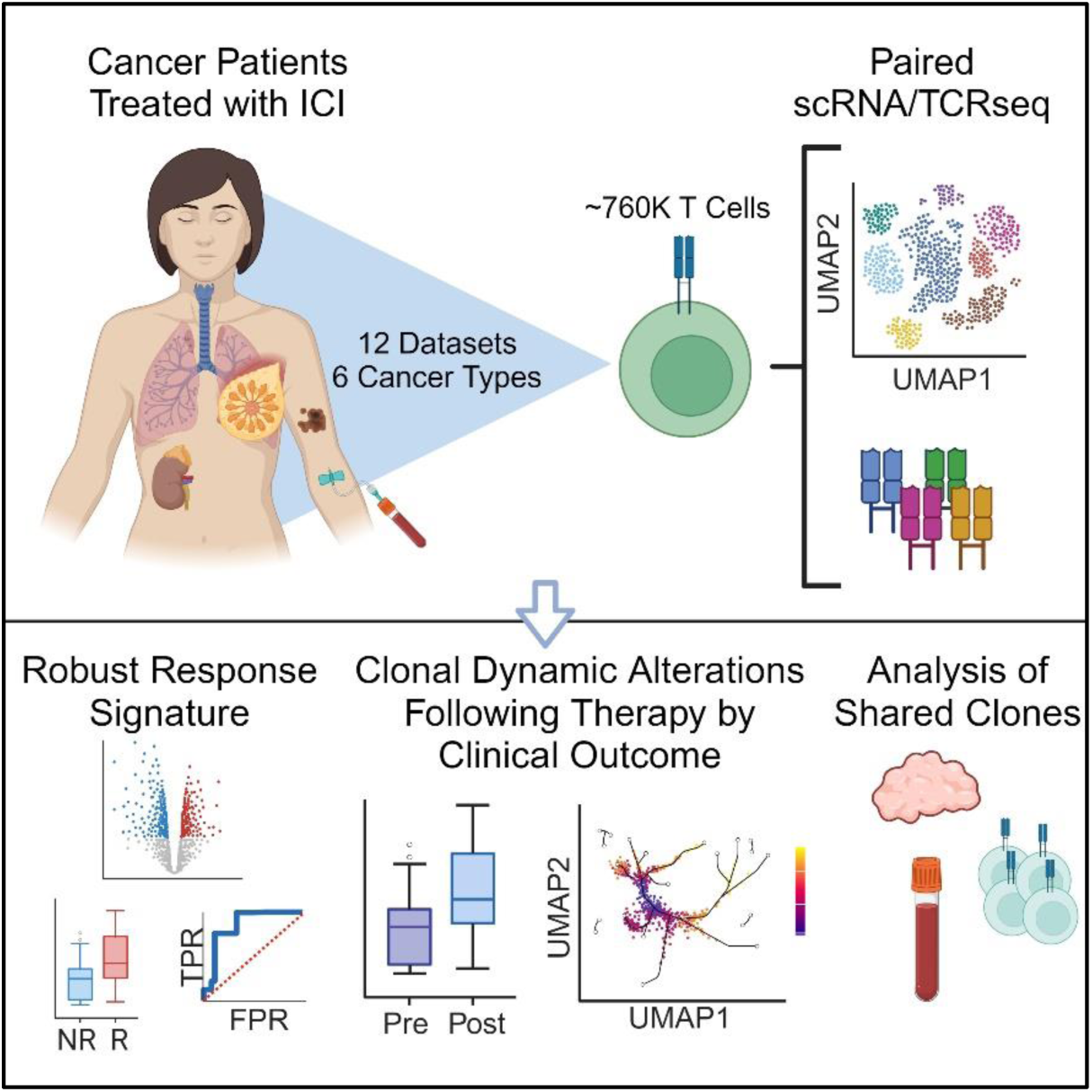

## Introduction

With the vast accumulation of single cell data during recent years, multiple studies utilized paired single-cell RNA/T-cell receptor sequencing datasets (scRNA/TCRseq) and placed T cell responses as a key component in the anti-tumor activity of cancer patients undergoing treatment with immune checkpoint inhibitors (ICI)^1^. Indeed, the mechanism by which T cells drive their anti-tumor activity, either by recruitment of novel clones from the blood into the tumor following ICI therapy (termed ‘T-cell clonal replacement’)^2^ or by stimulation of pre-existing intra-tumoral T cells (termed ‘T-cell clonal revival’)^3^, is still under debate^1,4^. For example, Yost et al. performed paired scRNA/TCRseq on longitudinal biopsies of basal and squamous cell carcinoma patients treated with ICI, and showed that expanded clones consisted of novel clonotypes that were not previously observed in the same tumor, indicating the limited reinvigoration capacity of pre-existing T cells in these tumors^2^. In contrast, Liu et al. utilized paired scRNA/TCRseq of NSCLC patients receiving ICI, showing that T cells from both periphery and local expansion within the tumor, replenish the pool of T cells with both new and pre-existing clonotypes^3^. Similarly, several studies of HNSCC patients, showed that 50-60% of on-treatment expanded clones were detected within tumors prior to therapy, and the rest were identified only in post rather than pre-treatment tumors^5,6^. The existence of both phenomena was also observed by van der Leun et al.^7^. Considering patient response, Au et al. similarly observed both novel and maintenance of pre-existing clones in post-treatment samples of renal cell carcinoma patients, and showed that only the latter was correlated with ICI response^8^. However, additional studies showed that the former is correlated with improved antitumor response^9–13^.

More recently, the phenotypic landscape of expanded clones was studied in the context of clinical outcome for ICI treated patients. Both Au et al. and Liu et al. showed upregulation of *GZMK* in expanded clones from responsive RCC and NSCLC tumors, respectively^3,8^. Shiao et al. similarly showed GZMK^+^ subpopulation having increased expansion following ICI therapy in triple-negative breast cancer (TNBC) patients, with higher expansion in patients responding to therapy compared to non-responders^14^. Zhang et al. showed that CD8-CXCL13 and CD4-CXCL13 cells are expanded in responsive tumors of TNBC patients, and demonstrated that responsive tumors exhibit higher levels of baseline CD8-CXCL13 and CD4-CXCL13 cells^15^. *CXCL13* was also found to be upregulated in expanded T cells from tumor samples of breast cancer patients receiving ICI^16^, and in tumor-enriched CD8^+^ and CD4^+^ clones of NSCLC patients receiving ICI^17^. However, identifying a signature for ICI response that will be robust across different studies spanning multiple cancer types is still challenging. Integrating and utilizing the massive amounts of existing scRNA/TCRseq data from multiple studies, can therefore offer sufficient statistical power for identifying robust transcriptional signatures in the context of clonal expansion. This, in turn, can provide valuable insights into the mechanism of patient response to therapy.

To address this challenge, we performed a comprehensive meta-analysis of paired scRNA/TCRseq data from 163 ICI treated patients across 6 cancer types, collected from 12 single-cell studies^2,3,5,8,14–19^. We profiled expanded clones from both tumor and blood samples using 767,606 single cells passing a strict quality control. We found that expanded clones can be in various cellular states, and are abundant in both responders and non-responders. Utilizing our integrated cohort, we investigated transcriptional changes in expanded CD8^+^ clones and revealed a robust gene signature that can significantly differentiate between responding and non-responding patients. We found this signature to be predictive in independent single-cell datasets of different cancer types and in sorted bulk samples. Moreover, analysis of persistent clones, as well as de-novo or dying clones enabled us to identify transcriptional changes that are differentially affected by treatment in responding or non-responding patients. Lastly, analysis of clones shared between tumor and blood samples resulted with distinct genetic programs that may be associated with clonal replacement or revival mechanisms, and are further linked to clinical outcome. The paired scRNA/TCRseq datasets used for this study can be easily accessed using our online repository (Single-Cell Vault) at: https://singlecellvault.net.technion.ac.il/, and are ready to be used by the scientific community.

## Results

### Profiling clonally expanded T cells in tumor and blood samples of ICI treated patients

To study clonal T cell signatures in tumor and blood samples of ICI treated patients, we analyzed 12 publicly available paired scRNA/TCRseq datasets consisting of 370 tumor samples and 90 blood samples taken from 163 ICI treated cancer patients across 6 cancer types^2,3,5,8,14–19^ (Table S1, Figure 1A-B). 767,606 single cells passed a strict quality control (QC) done separately on each of the two data modalities, scRNAseq and scTCRseq, and were used for further analysis (Method details). In addition, 12,407 genes shared between all datasets passed the QC process and were used for a further integration of all datasets while addressing batch effects between samples (Method details). Markov Affinity-based Graph Imputation of Cells (MAGIC)^20^ was first applied in order to detect possible drop-outs of *CD8A*/*B* or *CD4* (Method details, Supplementary Figure S1A-C). Following, single cells were labeled according to their membership in expanded or non-expanded T cell clones, using CDR3 sequence identity. Their clone size per sample was then quantified accordingly (Figure 1C, E, Method details). Finally, we further annotated the epitope of each single cell based on the CDR3 amino acid sequence using VDJdb^21^ as a reference database for epitope identity (Supplementary Figures S2A, S3A, Method details).

**Figure 1.**
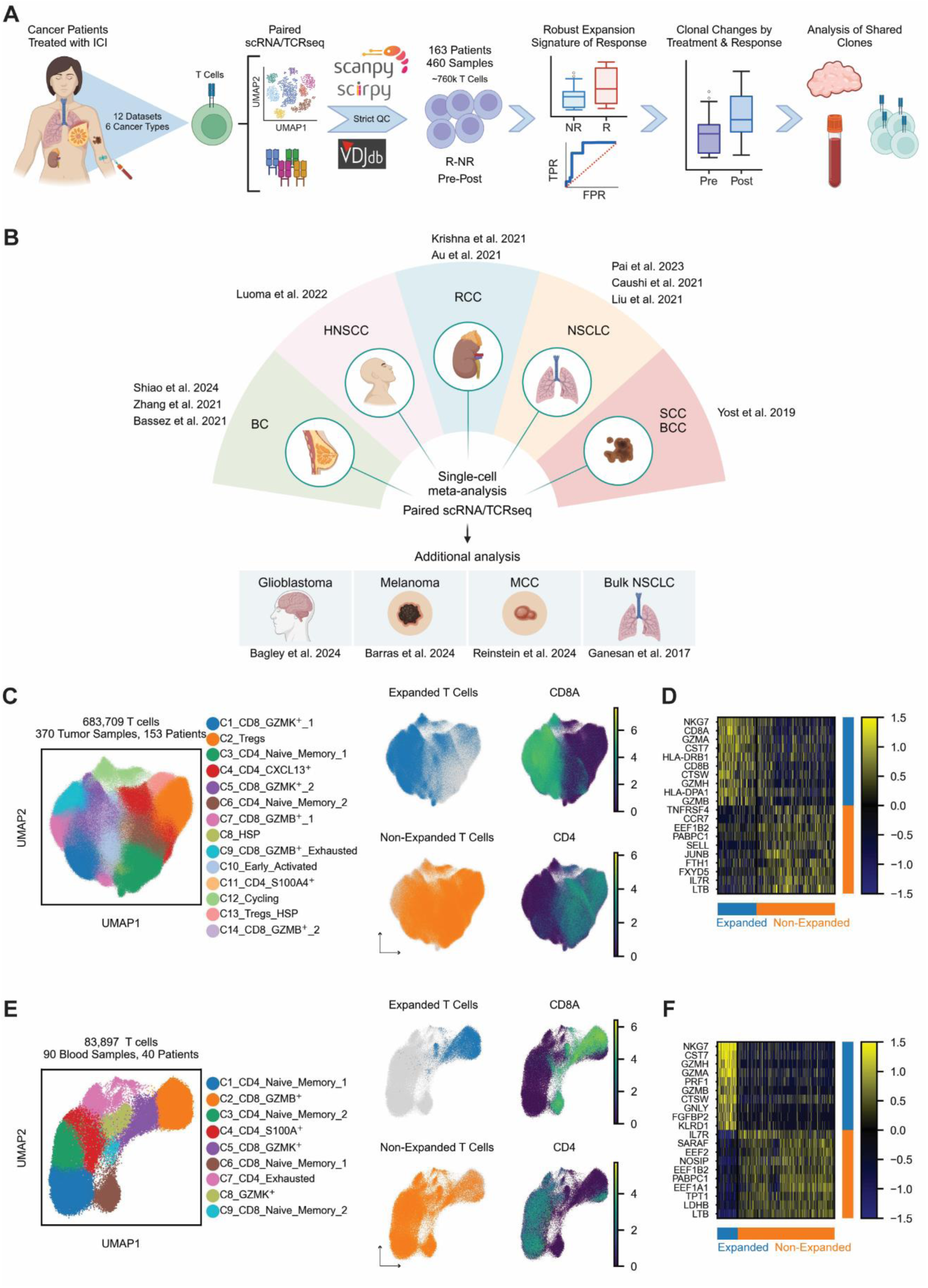
Profiling clonally expanded T cells in tumor and blood samples of ICI treated patients. A. A schematic workflow of the study. B. Schematic diagram showing all the utilized single-cell studies and cancer types. C. UMAP plot of 683,709 T cells from tumor samples having paired scRNA/TCRseq (left), and annotations for clonal expansion as well as expression of *CD8A* and *CD4* (right). D. Heatmap showing the top 10 differentially expressed genes between expanded and non-expanded T cells from tumor samples. Z-score was applied on the gene expression for visualization purposes, and only genes expressed in more than 10% of the cells in at least one of the groups were considered. E. UMAP plot of 83,897 T cells from blood samples having paired scRNA/TCRseq (left), and annotations for clonal expansion as well as expression of *CD8A* and *CD4* (right). F. Heatmap showing the top 10 differentially expressed genes between expanded and non-expanded T cells from blood samples. Z-score was applied on the gene expression for visualization purposes, and only genes expressed in more than 10% of the cells in at least one of the groups were considered. See also Figures S1-5, and Tables S1-2.

Differential gene expression between expanded and non-expanded T cells in both blood and tumor samples, showed that expanded T cells are predominantly CD8^+^ having increased expression of cytotoxicity and T cell activation markers (*NKG7*, *PRF1*, *GNLY*, *GZMA/B/H/K/M*), exhaustion markers (*LAG3*, *TIGIT*, *PDCD1*), as well as MHC-Class II genes (*HLA-DRB1/DPA1/DPB1/DRA/DQA1*/*DRB5*). Non-expanded T cells showed increased expression of naive-memory markers (*TCF7*, *IL7R*, *CCR7*, *SELL*, *LEF1*) as well as immune-regulatory markers such as *FOXP3*. Unlike blood samples, expanded T cells in tumor samples showed increased expression of genes such as *CXCL13* compared to non-expanded T cells (Figure 1D, F, Table S2).

To characterize the T cell clonality landscape and its association with clinical outcome in an unbiased manner, we conducted an unsupervised clustering analysis using the Leiden algorithm^22^ on single cells obtained from tumor and blood samples, yielding 14 and 9 T cell clusters respectively (Figure 1C, E, Table S2). Importantly, most clusters spanned evenly across the different datasets, and included different response status as well as varying levels of the expansion phenotype (Supplementary Figures S2B, S3B). Out of the 14 clusters obtained from tumor samples, six clusters contained a majority (>50%) of expanded T cells (Supplementary Figure S2B). Two of these clusters were GZMK^+^ (C1, C5), three were GZMB^+^ (C7, C9, C14), and one was a cluster of cycling T cells (C12). Examining differences in cluster abundance between responding and non-responding patients, we found an enrichment of C10 - Early_Activated in responders (p = 0.036, Supplementary Figure S2C), and significant enrichment of four clusters in non-responders (C3 – CD4_Naive_Memory_1, p = 0.021; C7 – CD8_GZMB^+^_1, p = 1.01*10^-4^; C11 – CD4_S100A4^+^, p = 1.01*10^-4^; and C14 – CD8_GZMB^+^_2, p = 0.013; Supplementary Figure S2C). Of these, only C7 and C14 (GZMB^+^) had a majority of expanded cells, demonstrating that T cell expansion on its own is not indicative of effective response. For blood samples, one cluster contained a majority of expanded T cells (C2, CD8_GZMB^+^), with more than 75% of the cells labeled as expanded (Figure 1E, Supplementary Figure S3B). However, none of the clusters were found to be significantly associated with clinical outcome (Supplementary Figure S3C).

To further examine differences in clonal expansion and their association with patient response, we focused solely on cells that are part of expanded clones in tumor samples, and compared their gene expression between responding and non-responding patients. As the abundance of expanded CD8^+^ clones per sample was significantly higher than that of CD4^+^ clones for both tumor and blood samples (p = 1.22*10^-61^ and 1.07*10^-25^ respectively, Supplementary Figure S1D), we decided to focus solely on expanded CD8^+^ clones for the following downstream analysis, while considering only clones that do not target any known non-cancerous antigens (method details). We found that responders show a significantly high expression of genes such as *GZMK*, *CXCR4*, *CXCL13*, and MHC-class II related genes - *HLA-DQA1*/*DQA2/DQB1*/*DRB5* (Table S2). Specifically, *CXCL13* was previously analyzed in a single-cell meta-analysis across five cancer types, and was found to be correlated with favorable response to ICI treatment^23^. Additional meta-analysis showed *CXCL13* to be a strong predictor of ICI response^24^. It was also shown to be exclusively expressed by intra-tumoral MCPyV-specific CD8^+^ T cells in MCC patients^25^, and by neoantigen-specific TCR clonotypes^26^, further suggesting its accurate indication for T cell specificity within tumors^25^. *GZMK* was shown to be abundant in nivolumab-bound expanded CD8^+^ T cells in responding patients having renal cell carcinoma^8^, and CD8^+^ GZMK^+^ cells were found to be significantly more abundant in acute myeloid leukemia patients responding to ICI-based therapy compared to non-responders^27^. In addition, *HLA-DQA1*/*DQA2* were recently shown as part of a predictive MHC-II signature for patient response, though in circulating T cells of ICI treated CRC and HNSCC patients^5,28^.

Non-responders on the other hand showed elevated expression levels of genes such as *CD52*, *S100A4*, *IL32*, *GZMB* and *ZNF683* (Table S2). Notably, antigen-activated T cells with high expression of *CD52* were previously shown to suppress other T cells^29^. *S100A4* was previously reported to be highly expressed in Tregs and exhausted T cells in gliomas, and was significantly associated with poor prognosis in glioma and glioblastoma patients^30^. *GZMB* was previously shown to be highly expressed in exhausted CD8^+^ T cells abundant in non-responding melanoma patients^31^, and in exhausted CD8^+^ T cells that characterized an exhausted tumor microenvironment (TME) for breast cancer patients^32^. Here, we emphasize the association between these genes and patient response within expanded CD8^+^ T cell clones, offering additional context for their involvement in treatment outcomes. This underscores their potential as biomarkers for predicting both favorable and unfavorable responses to ICI therapy across diverse cancer types.

A similar analysis with blood samples, showed that expanded CD8^+^ T cells in responders had a significant expression of genes such as *FOS*, *JUN* and *CX3CR1*, while non-responders had a significant expression of genes such as *EEF1G*, *DUSP2*, *KLF2* and *ZFP36L2* (Table S2). Indeed, *CX3CR1* was previously addressed as a blood-based biomarker of response to ICI treatment associated with good prognosis^33^.

Performing a pathway enrichment analysis using up-regulated genes in responders and non-responders of the integrated dataset, we found a significant upregulation of genes related to oxidative phosphorylation (OXPHOS) in non-responders for both tumor and blood samples (p = 9.21*10^-55^ and 0.0005 respectively), while responders showed higher TNF-alpha signaling (p = 3.91*10^-18^ and 0.005 respectively, Supplementary Figures S4-5, Table S2). This finding corresponds to a previous study showing that higher levels of OXPHOS in tumor and peripheral blood-derived CD8^+^ T cells correlate with ICI resistance^34^, and to our previous study devising an OXPHOS signature associated with poor response of ICI treated patients^35^. However, both studies used all single cells and were not focused on T cell clonal expansion. Overall, our analysis reveals distinct patterns of gene expression within expanded T cell clones in tumor and blood samples. While expanded T cells share a common activation and cytotoxic signature, this signature diverges significantly when comparing clones from responding and non-responding patients, highlighting the distinct molecular pathways that may drive differential therapeutic outcomes.

### Robust expansion signature differentiates between ICI responders and non-responders

To identify an expansion-related signature of response that will be robust across different cancer types, we conducted a differential expression analysis of single cells from expanded CD8^+^ clones between responders and non-responders as described above. To account for the variability across different datasets, we performed the analysis separately on each of the 9 different studies that contained both responding and non-responding samples^2,3,5,8,14,15,18,19^, out of the 12 studies that were analyzed in our integrated dataset (Method details, Figure 2A, Table S1, Table S3). We then applied a study-wise combined ranking of markers from all datasets (Method details) and obtained a ranked list of genes that are highly expressed in responders and non-responders across all datasets (Table S3). Following a robustness test (Method details), we resulted with a signature of 6 markers that are highly expressed in expanded clones from responders and 6 markers for non-responders (Figure 2B, Table S3).

**Figure 2.**
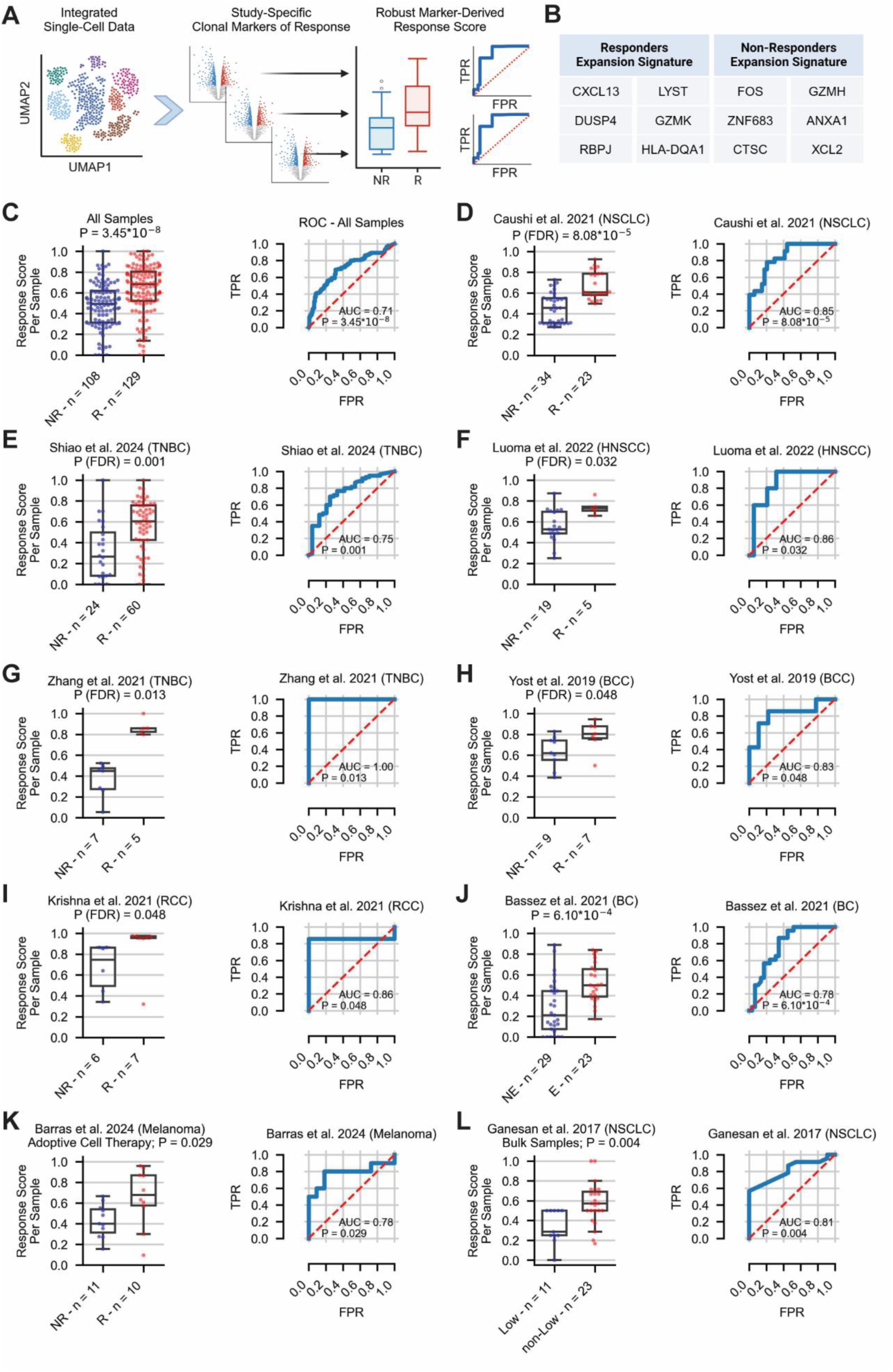
Robust expansion signature differentiates between ICI responders and non-responders. A. A schematic workflow for constructing the study-wise expansion-related response signature. B. Robust response signature of 6 markers obtained for responders (left) and non-responders (right). C. The performance of the response score across expanded T cells from 237 tumor samples spanning 9 single-cell studies^2,3,5,8,14,15,18,19^. ROC and the corresponding AUC achieved by the response score is shown on the right; Distribution of the response score in responders and non-responders is shown on the left. D-I. The performance of the response score across expanded T cells from tumor samples for selected datasets independently. J-L. The performance of the response score across expanded T cells from tumor samples of additional validation cohorts^16,41,42^. Abbreviations: R = Responders, NR = Non-responders, FPR = False positive rate, TPR = True positive rate. See also Figure S6, and Table S3.

The list of markers for responders include *CXCL13*, *DUSP4*, *RBPJ*, *LYST*, *GZMK* and *HLA-DQA1*. *CXCL13, GZMK* and *HLA-DQA1* were discussed earlier as they were significantly expressed in expanded clones of responders using the integrated dataset (Table S2). In addition, *DUSP4* was previously shown to be highly expressed in tumor-enriched CD8^+^ clones of metastatic NSCLC tumors^17^.

For non-responders, the obtained markers include *FOS*, *ZNF683*, *CTSC*, *GZMH*, *ANXA1* and *XCL2*. Upregulation of *FOS* expression in tumor infiltrating T cells was previously shown to promote tumor growth^36^ and *XCL2* was shown to be expressed in tumor-associated CD8^+^ T cells expressing exhaustion markers across four tumor types^37^. Of note, although *ZNF683* was previously associated with positive response to therapy in specific cancer types^38,39^, exploring its expression in multiple datasets clearly shows its up-regulation in non-responding patients.

Focusing on expanded CD8^+^ T cells, we scored each sample with a ‘response score’ based on both signatures, quantifying the ratio between expanded cells that express a more favorable response-related markers compared to markers associated with poor response. First, we found that this score significantly differentiates between responders and non-responders using all nine discovery datasets (p = 3.45*10^-8^, Area under the Curve [AUC] = 0.71, Figure 2C). Testing this signature for each single study separately, we managed, as expected, to significantly differentiate between responders and non-responders across multiple cancer types: for the NSCLC dataset of Caushi et al.^18^ we achieved a significant p-value of 8.08*10^-5^ and an AUC of 0.85 (Figure 2D). For the TNBC dataset of Shiao et al.^14^, we achieved p = 0.001 as well as AUC of 0.75 (Figure 2E). For the HNSCC dataset of Luoma et al.^5^, we achieved p = 0.032 as well as AUC of 0.86 (Figure 2F). For an additional TNBC^15^ dataset, as well as for BCC^2^ and RCC^19^ patients, we achieved p-values of 0.013, 0.048, and 0.048, respectively (Figure 2G-I). For 3 other datasets, this signature was insignificant (Supplementary Figure S6A-C). These include an additional SCC dataset^2^ showing the same trend with an AUC of 0.83, but lacks statistical power due to low sample size (Supplementary Figure S6A), and two other datasets of NSCLC and RCC patients^3,8^ (Supplementary Figure S6B-C). The latter results emphasize the ongoing challenge of finding reliable biomarkers of response that can be consistently generalized across different studies.

We next sought to validate our expansion-related response signature in additional datasets external to the discovery ones. Remarkably, in NSCLC patients having only non-responding patients^17^ the expansion signature was low and in a similar range to that of non-responders in other datasets (Supplementary Figure S6D). In two additional cohorts of breast cancer patients treated with ICI and having annotations for patient-level clonal expansion rather than clinical outcome^16^, our signature significantly differentiated between samples from both annotated groups, achieving a p-value of 6.10*10^-4^ and an AUC of 0.78 (Figure 2J). In de-novo glioblastoma patients treated with a combination of ICI and CAR therapy, all failed to respond^40^, our signature again achieved a similar low range of response score as was similarly demonstrated for non-responders in other datasets (Supplementary Figure S6D). In addition, we challenged our signature and examined its predictive power in a dataset of melanoma patients treated with adoptive cell therapy (ACT), having both responders and non-responders^41^. Notably, it significantly differentiated between the two groups, achieving a p-value of 0.029 and an AUC of 0.78 (Figure 2K). Finally, as our response signature was T cell related, we applied the response score on bulk samples sorted for T cells, of treatment-naive NSCLC patients^42^. Each sample was originally classified as TIL^hi^, TIL^int^ or TIL^lo^ according to the average number of CD8^+^ T cells that infiltrated the tumors^42^. Notably, our signature differentiated between samples classified as TIL^lo^ and those that were not TIL^lo^ (p = 0.004, AUC = 0.81, Figure 2L, Supplementary Figure S6E). This result, together with the improved survival seen for samples with higher density of CD8^+^ T cells^42^, further shows the potential of our signature to be predictive using sorted bulk samples, though more sorted data has to be collected in the context of response for treated patients. Overall, our study-wise signature of expansion-related response markers was able to robustly differentiate between responding and non-responding patients from multiple studies and across different cancer types.

### Transcriptional changes within persistent clones and their association with clinical outcome

To further delineate the transcriptional landscape of T cell clones, we applied consensus Non-negative Matrix Factorization (cNMF)^43^, a soft clustering approach that identifies gene programs and assigns each cell a program activity level between 0 and 1 (Method details). Our analysis identified 12 different programs across all tumor and blood samples (Figure 3A, Table S4). Nine of them were activity programs spanning different cellular types and states. Each program was annotated based on its top-ranked genes. Four out of the nine programs did not have clear annotations, but were found to be enriched with multiple cellular pathways, including TNF-alpha and mTOR signaling (Table S4, Method details).

**Figure 3.**
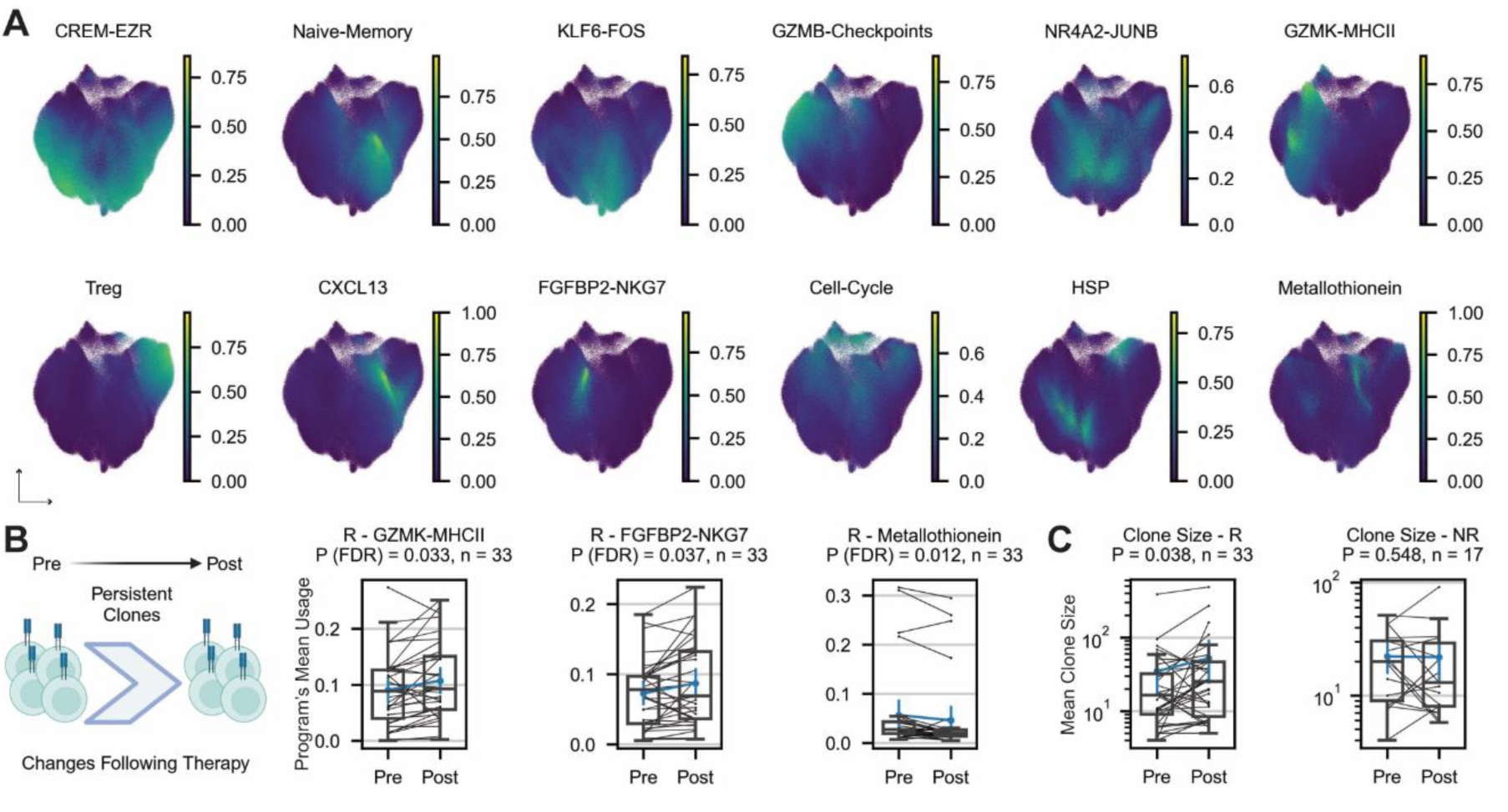
Transcriptional changes within persistent clones and their association with clinical outcome. A. Twelve transcriptional programs obtained using cNMF^43^. B. Changes following therapy of selected transcriptional programs for the top 5 expanded persistent clones per patient in responders (n = 33 patients). C. Change following therapy of the mean clone size for the top 5 expanded persistent clones per patient in responders (left, n = 33 patients) and non-responders (right, n = 17 patients). Abbreviations: R = Responders, NR = Non-responders. See also Figures S7-10, and Table S4.

Following, we examined how different gene programs of persistent CD8^+^ T cell clones, defined as clones found both at baseline and post-therapy samples, are affected by treatment in responding and non-responding patients. Across all datasets used in this study, 33 responders and 17 non-responders had tumor samples both at baseline and following administration of therapy, containing expanded CD8^+^ clones that persisted between both time-points. For each patient, we focused on the top 5 expanded clones, and measured how the activity level of each gene program is changing following therapy (Method details). In responders, we observed an increase in the activity of the cytotoxic programs GZMK-MHCII (p = 0.033) and FGFBP2-NKG7 (p = 0.037), with a decrease of the Metallothionein program (p = 0.012, Figure 3B). Metallothioneins are a family of metal-binding proteins known to down-regulate oxidative stress^44^, commonly elevated in cancer. Their down-regulation in responders may point to effective tumor clearance. In addition, responders demonstrated a significant increase in the mean clone size following treatment (p = 0.038, Figure 3C). In contrast, non-responders did not demonstrate any significant change in gene programs following therapy (Supplementary Figure S7A-B), or a change in clone size (p = 0.548, Figure 3C). These findings reinforce the enhanced intra-tumoral reinvigoration capacity of T cell clones in responders, as previously noted^3^. Moreover, in non-responders, the lack of changes in key transcriptional programs may be attributed to factors such as pre-existing immune dysfunction, a suppressive TME, intrinsic resistance mechanisms, or impaired immune activation, preventing effective engagement of the immune response following ICI therapy.

Searching for transcriptional changes in blood samples of 7 responding and 21 non-responding patients, we did not identify any significant changes in program activity following therapy, or in clone size (Supplementary Figure S8A-D). This finding further demonstrates the importance of the TME in inducing transcriptional changes and highlights the challenge of identifying biomarkers for patient response based on blood samples.

As described earlier, oxidative phosphorylation was found to be highly upregulated in expanded clones in non-responders for both tumor and blood samples (Supplementary Figures S4-5). Following this finding which marks the importance of metabolism in regulating T cell function, and our previous work showing a predictive metabolic sub-classification of T cells^35^, we performed an additional analysis focused solely on metabolic genes^45^ (Method details, Table S4). This process resulted with 6 different metabolic activity programs that were similarly tested for transcriptional changes following treatment (Supplementary Figure S9, Table S4). We found an increase of the LDHB-GSTK1 program following treatment in tumor samples of responders (p = 0.029, Supplementary Figure S9B). This metabolic program is not exclusively expressed in a specific T cell state but rather spans distinct ones, including naive-memory and effector T cells. Notably, we previously showed that top genes of this program (*LDHB*, *GSTK1*, *DGKA*, *APRT*, *MGAT4A* and *NMRK1*) are predictive of cancer patient response to ICI^35^. This program was also found to be highly abundant in blood samples, and is highly correlated with the Naive-Memory program described above (Supplementary Figures S9C, S10A).

Finally, we addressed transitions between gene programs by tracking the max activity of programs in individual persistent clones over time (Supplementary Figure S10B-C). We found that in responding patients, a subset of cells found in an exhaustion state as depicted by the GZMB-Checkpoints program, was able to transition into the cytotoxic GZMK-MHCII program. Notably, this transition was not observed in non-responders. Similar observation was previously reported for intrahepatic cholangiocarcinoma patients receiving combined therapy with ICI, where such a transition from CD8 GZMB^+^ to CD8 GZMK^+^ facilitated good response to therapy^46^. In addition, we found that a larger fraction of the CREM-EZR program switched to the cytotoxic GZMK-MHCII program in responding compared to non-responding patients. However, the role of this program has not been widely studied and showed ambiguous associations to different phenotypes of T cells such as exhausted^47^ and effector memory^48^. From a metabolic point of view, we observed a higher fraction of clones transitioning from the suppressive metabolic CHST12-CD38 program into the beneficial LDHB-GSTK1 program in responders compared to non-responders (Supplementary Figure S10B-C). Taken together, these results demonstrate again an extended reinvigoration ability of T cells following ICI treatment in responding patients.

### Pseudo-temporal changes of CD8^+^ expanded clones by clinical outcome

We next sought to analyze the pseudo-temporal dynamics of expanded clones at the single-clonal level. To this end, we analyzed expanded clones in a pseudobulk manner per sample using the mean expression of all single cells per each expanded clone (Method details, Figure 4A). This approach resulted with 7,945 expanded CD8^+^ clones from all tumor samples across all datasets, that were considered for further analysis. We then applied GeneTrajectory^49^ - an approach that identifies trajectories of genes rather than of cells and outperforms multiple cell-trajectory methods in recovering the gene order for both cyclic and linear processes^49^ (Method details). We found that the trajectory of CD8^+^ clones demonstrates the clonal transition from a naive-memory state (*CCR7*, *IL7R*, *TCF7*) towards activation (*GZMK*), clonal exhaustion (*CTLA4*, *PDCD1*, *HAVCR2*), reaching eventually to *CXCL13* expression and a cell-cycle state (*TYMS*, *TK1*, *MKI67*, *DHFR*, Figure 4A-B, Table S5).

**Figure 4.**
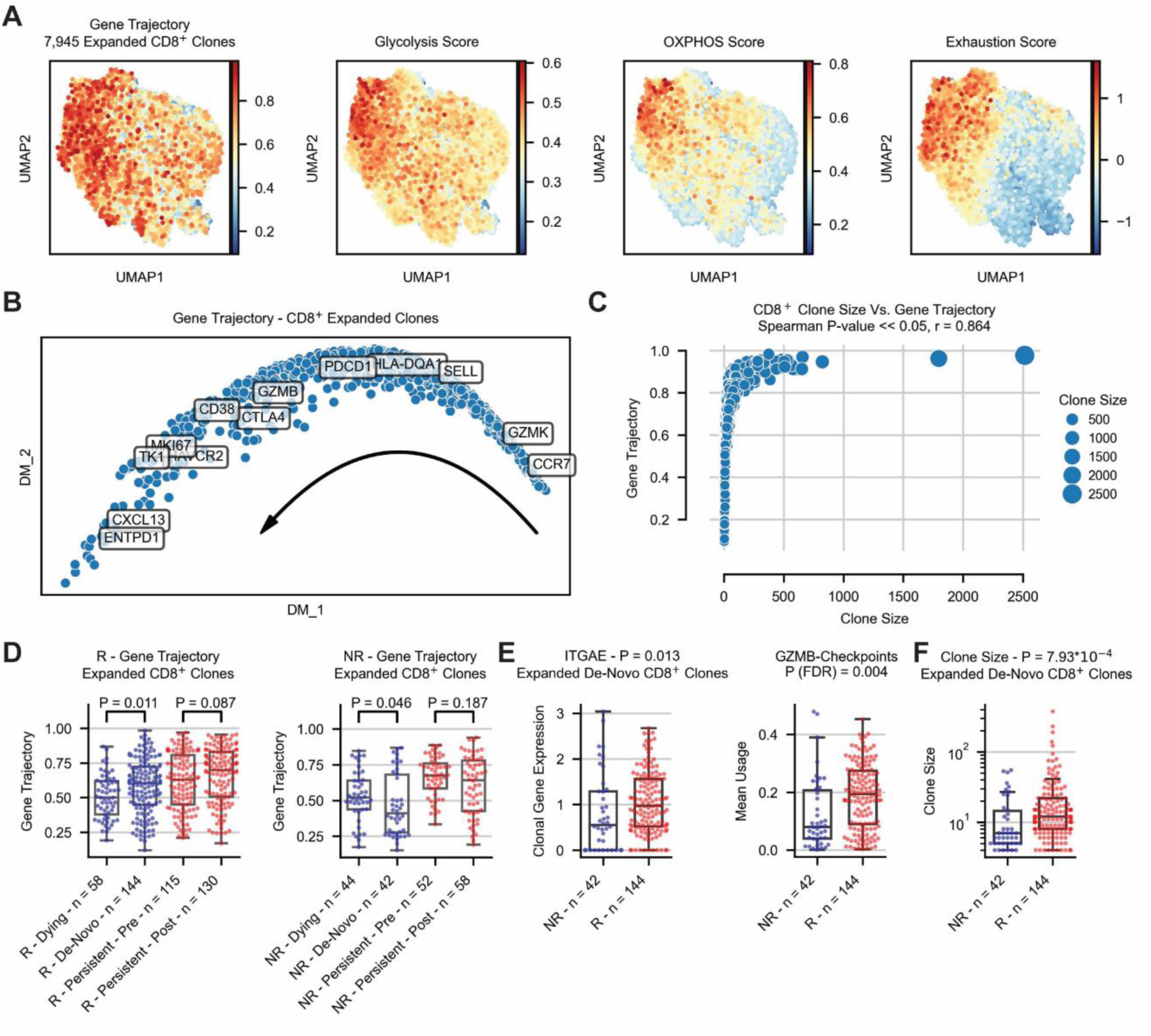
Pseudo-temporal changes of CD8^+^ expanded clones by clinical outcome. A. Reflection of the obtained gene trajectory^49^ over the UMAP plot of 7,945 expanded CD8^+^ clones from tumor samples, showing how the genes are expressed across different regions of the clonal embedding (left), as well as clonal scores of glycolysis, oxidative phosphorylation and exhaustion (right). B. Diffusion map visualization of the CD8^+^ clonal gene trajectory, including annotations for selected genes. C. Spearman correlation between the gene trajectory reflection of each CD8^+^ expanded clone and its clone size. D. Gene trajectory values for the top 5 expanded CD8^+^ clones per sample in responders (left) and non-responders (right), separated by dying, de-novo and persistent clones. E. Difference in *ITGAE* expression (left) and the activity of the GZMB-Checkpoints program (right) per clone, for the top expanded de-novo CD8^+^ clones in responders and non-responders. F. Difference of clone size for the top expanded de-novo clones per sample between responders and non-responders. Abbreviations: R = Responders, NR = Non-responders, OXPHOS = Oxidative phosphorylation. See also Figures S11-12, and Table S5.

Considering the importance of metabolism in T cell expansion, we further tested the Spearman correlation of the gene trajectory with 96 metabolic pathways mapped by the Recon 2 metabolic reconstruction^45^ (Method details). This analysis showed that Glycolysis and Oxidative phosphorylation are the top two metabolic pathways with a high activity score across all expanded CD8^+^ clones, and are also highly correlated with the gene trajectory (Method details, Figure 4A, Table S5). Each clone was also scored for exhaustion and memory signatures (Method details), demonstrating the clonal transition between these cellular states (Figure 4A, Supplementary Figure S11A). Notably, we also found that the size per sample of each expanded clone is correlated with the gene trajectory (p < 2.22*10^-308^, Figure 4C), suggesting that clonal states are also related to changes in clone size.

To examine how ICI therapy affects clonal development, we explored the position of different T cell clones along the trajectory. We focused on de-novo, dying and persistent clones, such that de-novo clones are defined as those expanded only following treatment, and dying as those expanded only at baseline. Analyzing the top 5 expanded CD8^+^ clones per sample in patients having longitudinal biopsies (Method details), we found significant differences in the gene trajectory score between dying and de-novo clones, such that de-novo clones in responders are found further down the trajectory path compared to dying clones, with an opposite trend in non-responders (Figure 4D). Such differences were not significant for persistent clones in either response groups.

To further explore this finding, we considered the activity of transcriptional programs along the gene-trajectory, and observed higher activity of the GZMB-Checkpoints program in the more advanced pole of the trajectory (Supplementary Figure S11B). Indeed, we found increased activity of this program in de-novo clones of responding patients (p = 0.004, Figure 4E, Supplementary Figure S12). This finding coincides with the significant difference in clone size of de-novo clones between both response groups (p = 7.93*10^-4^, Figure 4F), and with the correlation of clone size to the gene trajectory (Figure 4C). Interestingly, the GZMB-Checkpoints program is associated with elevated expression of *ITGAE*, a marker of tissue resident memory T cells, which ranked highest in this program, compared to all other transcriptional programs (Supplementary Figure S11B, Table S4). Indeed, a comparison of top expanded de-novo CD8^+^ clones between responders and non-responders demonstrated significantly higher *ITGAE* expression in responders (p = 0.013, Figure 4E). Collectively, these findings suggest that de-novo clones in responders are more likely to originate within the TME, whereas non-responders exhibit relatively lower levels of intra-tumoral emergence.

### Intra-tumoral CD8^+^ clones shared with blood are associated with non-responders

We next examined the relation between expanded clones shared by tumor and matched blood samples by utilizing three datasets containing 20 non-responding and 10 responding patients having tumor with matched blood samples^5,15,19^ (Figure 5A). Overall, out of 20 non-responders and 10 responders, a majority of 15 non-responders (75%) and only 4 responders (40%) had expanded CD8^+^ clones that were shared between both tissue types. However, when addressing transcriptional changes of shared clones between blood and tumor samples (Method details), similar changes in gene programs appeared regardless of clinical outcome. These changes include a decreased activity of multiple programs such as: FGFBP2-NKG7, Treg and Naive-Memory, with increased activity of several other programs such as: CREM-EZR, KLF6-FOS, GZMB-Checkpoints, NR4A2-JUNB, GZMK-MHCII and HSP (Supplementary Figure S13A-B). The reduced activity of the Naive-Memory program, as well as the higher activity of cytotoxic programs in tumor compared to blood samples, coincides with the loss of the bystander phenotype of peripheral T cells following their infiltration into the tumor, regardless of clinical outcome. This loss of bystander phenotype also corresponds to the changes seen in the activity of the metabolic LDHB-GSTK1 program which is highly correlated to the Naive-Memory program (Supplementary Figures S10A, S14).

**Figure 5.**
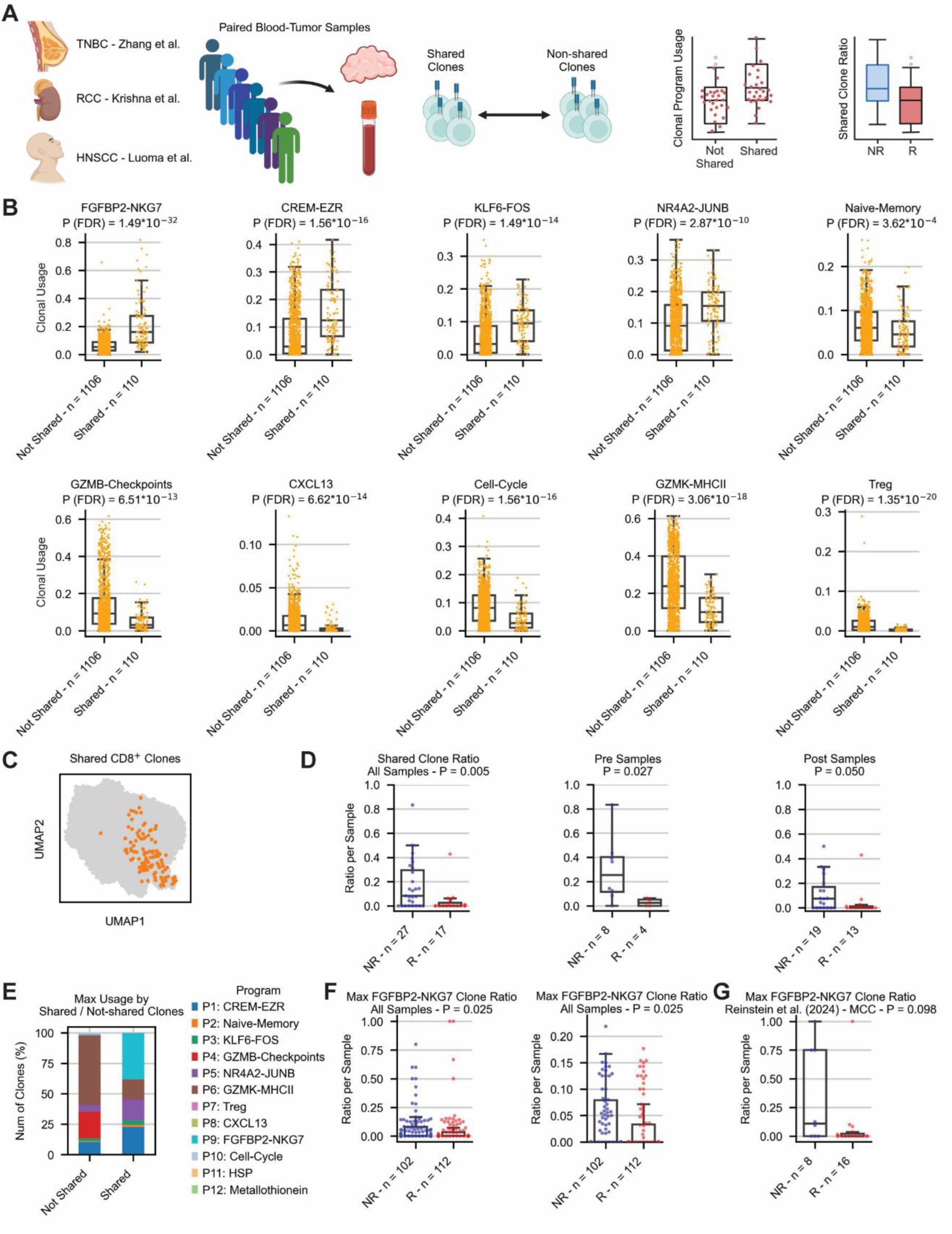
Intra-tumoral CD8^+^ clones shared with blood are associated with non-responders. A. A schematic workflow for the analysis of shared clones, using three single-cell studies^5,15,19^. B. Difference in the usage of selected transcriptional programs per clone for shared and non-shared clones. C. UMAP plot demonstrating the location of shared CD8^+^ clones on their clonal embedding. D. Abundance of shared clones out of all the expanded clones per sample, and its difference between responding and non-responding samples. Results are presented using pre and post samples together (left), as well as each time-point separately (right). E. Fraction of shared and non-shared clones based on the max activity of transcriptional programs per clone. F. Fraction of expanded clones having max usage of the FGFBP2-NKG7 program, out of all the expanded clones per sample, tested between responders and non-responders using the full range of ratios (left), or a zoomed-in range up to a ratio of 0.2 (right). G. Fraction of expanded clones having max usage of the FGFBP2-NKG7 program, out of all the expanded clones per sample, tested between responders and non-responders of MCC patients treated with ICI^50^. Abbreviations: R = Responders, NR = Non-responders. See also Figures S13-16.

Following the significant number of non-responding patients with clones shared between tumor and blood, we hypothesized that distinct transcriptional programs are active in such clones as compared to those that exist only in the tumor. To examine that, we labeled clones as “shared” or “non-shared” accordingly (Method details). This classification resulted with 110 shared clones and 1,106 non-shared clones that were expanded across 44 tumor samples from all 30 patients (Figure 5B-C). Examining the activity of our previously defined cNMF programs (Table S4), we observed a significant expression of four programs in shared clones, with the most significant expression of the FGFBP2-NKG7 program (p = 1.49*10^-32^, Figure 5B, Supplementary Figure S15A), and a significantly lower expression of *ITGAE* in shared clones compared to non-shared clones (p = 8.21*10^-10^, Supplementary Figure S15B). Calculating the ratio of shared clones out of all expanded CD8^+^ clones per sample, we found a significant abundance of shared clones in non-responders (p = 0.005, Figure 5D), further suggesting that non-responders show higher recruitment of clones from the blood, whereas responders demonstrate higher intra-tumoral reinvigoration. This significant difference between both response groups occurred regardless of treatment time-point (Figure 5D).

To examine the generalization of these findings in datasets that do not have paired blood samples, we searched for a signature that is abundant in shared clones, and has low abundance in non-shared clones. Indeed, while considering the max activity of each program, 38.18% of the shared clones had max activity of the FGFBP2-NKG7 program, compared to only 0.72% of the non-shared clones (Figure 5E). We therefore used this program as a potential signature for shared clones, and identified it in additional seven datasets having only tumor samples with annotations for clinical outcome^2,3,8,14,17,18^ (Methods details). Interestingly, clones with max activity of this program were significantly more abundant in non-responders (p = 0.025, Figure 5F), further suggesting that shared clones are more abundant in non-responders. These results were also validated in an additional cohort of MCC patients treated with checkpoint inhibitors^50^ (Method details, Supplementary Figure S16), showing higher fraction of clones with max activity of the FGFBP2-NKG7 program in non-responders (p = 0.098, Figure 5G). These findings coincide with a recent study of melanoma patients receiving adoptive cell therapy (ACT), showing that blood-borne clonotypes from ACT products repopulated mostly blood in responders, while infiltrating tumors of non-responders^51^. It is also reflected by the higher activity of the FGFBP2-NKG7 program in de-novo clones in non-responders compared to responders (p = 0.006, Supplementary Figure S12), further suggesting that de-novo clones in non-responders rather originate from blood-borne clones. Of note, these results stand in contrast to a previous study showing that T cells, especially in responsive patients, are replenished from sites outside the tumor^9^, and to a study of HCC patients showing higher abundance of shared clones in responders^52^. However, the latter computed the fraction of shared TCRs relative to the total number of TCRs per sample, while we computed this fraction using only expanded CD8^+^ clones. Taken together, our results highlight potential mechanism by which responders and non-responders reactivate or recruit their intra-tumoral expanded clones via clonal revival or clonal replacement, respectively.

## Discussion

Clonal expansion has been widely studied in the context of response to checkpoint immunotherapy. However, the small sample size of each individual study limits the scope and generalization of the drawn conclusions. Our meta-analysis aims to bridge this gap by analyzing multiple datasets, thus revealing signals that are otherwise hidden. As a result, our findings provide robust insights that can further shed light on mechanisms of response to therapy, and highlight meaningful biomarkers of response that have the potential for clinical applications.

Specifically, our study profiled genes that differentiate between expanded and non-expanded T cells for both tumor and blood samples, and tested for differences between expanded clones in responders and non-responders. Conducting a study-wise analysis, we devised a predictive gene signature of expanded CD8^+^ T cells that significantly differentiates between responders and non-responders across multiple datasets, spanning several cancer types. We validated this signature on additional cohorts of ICI treated patients and further showed its performance on sorted bulk samples of NSCLC patients. We then detected continuous genetic programs with different activity levels, and showed their changes following therapy in persistent clones. This analysis demonstrated substantial differences between responding and non-responding patients, highlighting increased reinvigoration capacity in responding patients. Using gene-trajectory analysis, we recovered the gene ordering of expanded CD8^+^ clones and showed the pseudo-temporal changes of their clonal states. We found that de-novo clones from responding patients are found further down the trajectory path, which together with their transcriptional activity suggested that these clones originate from within the TME, as opposed to non-responding patients. Finally, analyzing clones that were either shared or not shared between tumor and blood samples, we identified distinct gene programs that may underlie differences in T cell function, activated through either the replacement or revival processes.

The transcriptional profiling of T cell clones provides a nuanced understanding of the mechanisms driving response and resistance to ICI therapy. In responders, persistent clones exhibited upregulation of the beneficial GZMK-MHCII program. This indicates that these clones are not only persisting but also becoming more functionally active, contributing to effective anti-tumor responses. In non-responders, such clones did not show significant upregulation of programs post-therapy. This lack of transcriptional changes indicates that these clones may remain in a state of exhaustion or suppression, failing to contribute effectively to anti-tumor activity.

Furthermore, our findings highlight that intra-tumoral clones shared with blood are predominantly associated with non-responders to ICI therapy, which indicates potential systemic circulation of less effective T cells. The fact that the FGFBP2-NKG7 program majorly characterized shared clones abundant in non-responders (Figure 5B, D-G), but was also increased in persistent clones for responders (Figure 3B), possibly implies that both clonal replacement and revival are executed in responders and non-responders, but to a different extent. In addition, clones with high activity of this program were located at one pole of the clonal embedding (Supplementary Figure S11B), further implying of their distinct pseudotemporal state. These results underscore the importance of understanding the clonal architecture and dynamics within the TME and peripheral blood to better predict and improve responses to ICI therapy.

Interestingly, several genes from our identified signature that were associated with poor response, including *ZNF683*, were expressed in multiple transcriptional programs that were activated in different contexts. This includes the FGFBP2-NKG7 and the GZMB-Checkpoints programs that were significantly activated in blood-tumor shared and non-shared clones, respectively. Together with the fact that *ZNF683* was previously linked to improved response in ICI-treated Richter syndrome patients^38^, and to revival of pre-existing *ZNF683^+^*clones in HNSCC patients responding to ICI treatment^39^, this finding emphasizes the importance of the contextual co-expression of genes in relation to clinical outcome, as well as differences between gene expression in different cancer types and tissues.

Taken together, our analysis underscores the importance of T cell clonal expansion and specific genetic programs in determining patient response to ICI therapy. Specifically, it suggests that clonal revival dominates in responders, and clonal replacement is more common in non-responders, highlighting the critical role of reversible T cell exhaustion and the ability of existing T cell clones to be reactivated in successful therapy. This would suggest that reviving existing tumor-infiltrating lymphocytes is a more reliable mechanism of anti-tumor activity than recruiting new clones from the blood, at least in the context of ICI therapy. However, more data containing longitudinal biopsies with matched blood samples is required in order to better quantify this phenomenon in both response groups.

## Limitations of the study

While a broad meta-analysis of paired scRNA/TCRseq data can provide important insights on clonal expansion for ICI treated patients, this approach requires dissociation of cells from their tissue, resulting in loss of spatial information and the relative positioning of cells which is critical to determine neighboring structures, cellular subtypes and cellular states^53^. Therefore, such meta-analysis should also be studied in the locational context using spatial transcriptomics, when more data become available. In addition, our results emphasize the ongoing challenge of finding a robust signature of biomarkers that is predictive of patient response across different studies and cancer types. Despite its high performance, our devised signature failed to differentiate between responders and non-responders in two datasets^3,8^ (Supplementary Figure S6B-C), while succeeding in other datasets of the same cancer type (Figure 2). Such discrepancies between studies of the same cancer type are a result of multiple confounding factors unequally controlled in different studies, including treatment regimen, sampling time, and tumor characteristics^1^. One specific concern is the treatment factor. While all patients in this study received ICI-based therapy, the data represents a heterogeneous group with variations in the specific drugs used and their combination with other treatments, such as chemotherapy, radiation, CAR therapy, or other ICIs. This variation in treatment regimens among patients likely influence the transcriptional signatures observed. However, due to the large sample size we were still able to identify common signatures underlying ICI-based treatment. Moreover, our analysis was able to identify signatures that are common across cancer types, but more data is needed in order to address cancer-type-specific signatures. Another crucial point to note is validation in bulk samples. Since our signature is T-cell based and specifically related to expanded T cells, validating it in bulk RNAseq remains a significant challenge. Lastly, functional studies are required in order to elucidate the causal effects of clonal states described in this study on cancer patient response to checkpoint immunotherapy.

## Author Contributions

O.S and K.Y conceived the idea. O.S and K.Y designed the study. O.S performed the analysis. A.P created the manuscript-related website. O.S and K.Y wrote the manuscript.

## Methods

### Resource availability Lead contact

Further information and requests for resources should be directed to and will be fulfilled by the lead contact, Keren Yizhak.

### Materials availability

This study did not generate new unique reagents.

### Data and code availability

- This paper analyzes existing, publicly available data. The accession numbers for these datasets are summarized in Table S1. Paired scRNA/TCRseq data used for the integrated analysis are accessible following our quality-control at: https://singlecellvault.net.technion.ac.il/
- This paper does not report original code.
- Any additional information required to reanalyze the data reported in this paper is available from the lead contact upon request.

## Method details

### Paired scRNA/TCRseq datasets and preprocessing

12 single-cell datasets of ICI-based treated patients having paired scRNA/TCRseq were collected together with their annotations for clinical outcome and treatment time-point^2,3,5,8,14–19^. Nine datasets contained tumor biopsies while three datasets contained both tumor and blood samples (Table S1).

All scRNAseq datasets were droplet-based and contained read counts. For each dataset, we followed similar preprocessing steps to those conducted by Tang et al.^54^ using Scanpy^55^: We removed cells expressing less than 200 genes and removed genes that were expressed in less than 3 cells. We also removed cells having more than 10% of mitochondrial gene-count. In the dataset of Krishna et al.^19^, scRNAseq was provided following the authors’ original QC while excluding mitochondrial genes from the provided gene expression matrix. In that case only, we used the expression matrix as provided by the authors following their QC, retaining cells with less than 20% of mitochondrial gene-count. We also applied Scrublet^56^ to remove potential doublets and filtered-out cells having *‘doublet_score’* larger than 0.3.

The clinical metadata for each dataset was used as provided by the authors. In the dataset of Luoma et al.^5^, response was provided both by RECIST^57^ and as pathological response. In that case, and because many patients had a “not measurable” response according to RECIST but did have pathological response, we considered responders as those with high pathological response and non-responders with a non-high pathological response. For the dataset of Caushi et al.^18^ we considered samples with major pathological response (MPR) as responders and those with non-MPR as non-responders. For the dataset of Shiao et al.^14^ we considered samples with complete pathological response (pCR) as responders, and those with non-pCR as non-responders. For the dataset of Krishna et al.^19^, we annotated the response status of each patient according to the original description of each individual patient provided by the authors. For datasets providing response according to RECIST^15,17^, we labeled samples with complete/partial response (CR/PR) as responders and those with stable/progressive disease (SD/PD) as non-responders, as was similarly done previously^31,35^. For the datasets of Yost et al.^2^, Liu et al.^3^, and Au et al.^8^, we annotated each sample according to the original annotations provided by the authors. The two breast cancer cohorts of Bassez et al.^16^ were not provided with labels of clinical outcome, but rather with a patient-level annotations for clonal expansion that were used as provided by the authors.

For the scTCRseq datasets, we rearranged each dataset to be compatible with the Adaptive Immune Receptor Repertoire (AIRR) schema that was further preprocessed by Scirpy^58^ and could be directly uploaded using the *‘scirpy.io.read_10x_vdj’* function. For each scTCRseq dataset, we applied quality control based on the standard protocol suggested by the authors. In short, we removed cells having multi-chains, orphan VJ or orphan VDJ chains. We also removed a single cell having “ambiguous” receptor type, indicating it has both BCR and TCR.

### Integration of scRNA/TCRseq datasets

Following the quality control process described above, we integrated all processed datasets resulting with a total of 683,709 single cells from tumor samples and 83,897 single cells from blood samples, all with paired scRNA/TCRseq. Overall, 12,407 genes existed across all datasets and were used for further analysis, such that tumor and blood samples were analyzed separately.

For the integrated scRNAseq datasets, we normalized the expression level to a standard target sum of 10,000 reads per cell and then applied log2-transformation. Highly variable genes were calculated using *‘scanpy.pp.highly_variable_genes’* with a *‘batch_key’* of *‘cancer type’* in order to preserve biological differences between the different cancer types analyzed in this study. We then calculated a 40-component PCA and applied Batch Balanced K-Nearest Neighbors (BBKNN)^59^ with a ‘*batch_key’* of *‘sample’* in order to remove batch effects between samples. UMAP^60^ was used for dimensionality reduction and data visualization.

### Markov Affinity-based Graph Imputation of Cells (MAGIC) for detection of drop-outs and labeling of expanded clones

In order to identify possible drop-outs in our integrated dataset, we applied MAGIC^20^ on the log2-normalized count matrix using all the genes and single-cells passing our QC. We used the default arguments of the algorithm’s implementation in Scanpy^55^. We first fitted a density curve to the log2-transformed imputed gene expression of *CD8A/B* and *CD4*, and set the expression threshold for each gene as the trough of the bimodal density curve (Supplementary Figure S1A-B). For non-imputed gene expression, log2-transformed expression threshold was set as 1. Overall, all the T cells that passed our QC were further divided into four subtypes (Supplementary Figure S1C): we considered a single cell to be CD8^+^, when the imputed or non-imputed gene expression of either *CD8A* or *CD8B* was above the threshold. Similarly, a single cell was considered to be CD4^+^, when the imputed or non-imputed gene expression of *CD4* was above the threshold. We also defined double-positive cells as those that were considered to be both CD8^+^ and CD4^+^. Double-negative cells were those that were both CD8^-^ and CD4^-^.

We then defined clonotypes based on the identity of the CDR3 nucleic acid sequence of each cell using *‘scirpy.tl.define_clonotypes’*, such that both the VJ and VDJ CDR3 sequences had to match and the T cell subtype of all the cells in each clone is the same (i.e. CD8^+^, CD4^+^, double-positive or double negative). In cases where more than one pair of VJ and VDJ sequences was detected per cell, we considered the most abundant pair of VJ/VDJ chains where applicable. In the dataset of Krishna et al.^19^, scTCRseq was provided only with the amino acid sequence of the CDR3 region. In that case only, clonotypes were defined based on the amino acid identity of the CDR3 sequence.

We defined a clonotype as “expanded” per sample according to Shiao et al.^14^, such that “expanded clones” were those that contained more than 1.5x the median number of cells found in each clonotype in that sample, while excluding singletons. In order to conduct further analysis at the level of each sample, we assigned each clone with a unique ID such that the same clone appearing in multiple samples from the same patient received different ID, indicating the clone’s sample of origin. We then used VDJdb^21^ as a reference database for annotating epitopes based on amino acid sequence identity according to the standard protocol suggested by the authors^58^ (Supplementary Figures S2A, S3A).

### Clustering and differential gene expression analysis

Unsupervised clustering was done with the graph-based Leiden^22^ algorithm as implemented by Scanpy. Differentially expressed genes (DEGs) per cluster were calculated using the *‘scanpy.tl.rank_genes_groups’* function with the Wilcoxon rank-sum test (Table S2), followed by manual annotations for cluster-names according to the top differentially expressed genes of each cluster. Similarly, we used the Wilcoxon rank-sum test to calculate differentially expressed genes between expanded and non-expanded cells (Table S2).

To identify clusters that are significantly associated with patient response, we calculated for every cluster the fraction of cells assigned to that cluster in every sample. We then conducted a two-sided Wilcoxon rank sum test between these fractions in responders and non-responders. This process was done for each cluster and was corrected for multiple hypothesis using the Benjamini-Hochberg false discovery rate^61^. Adjusted p-values < 0.05 were considered as significant (Supplementary Figures S2C, S3C).

### Extraction of CD8^+^ T cells from expanded clones that do not target known non-cancerous antigens

As the abundance of expanded CD8^+^ clones per sample was significantly higher than all of the other clonal subtypes (Supplementary Figure S1D), we focused solely on CD8^+^ T cells for the entire downstream analysis. In addition, we considered only clones that do not target a known non-cancerous antigen based on the VDJdb^21^ annotations that we assigned for every single-cell as described above. This enabled us to focus only on expanded CD8^+^ clones that potentially target cancerous neoantigens. In cases where cells of the same clone could also target a known antigen due to an extra VJ or VDJ chain, or due to multiple full VJ/VDJ pairs, we considered only those where the majority of cells from the same clone did not target any known non-cancerous antigen. We then used the Wilcoxon rank-sum test to calculate differentially expressed genes between expanded CD8^+^ T cells from responders and non-responders, as described above (Table S2).

### Analysis of upregulated genes in expanded CD8^+^ T cells by response

The GSEApy enrichr module^62^ was used to conduct over-representation analysis between expanded CD8^+^ T cells in responders and non-responders. For each of these groups, we extracted its upregulated genes having adjusted p-value < 0.05, as well as positive and negative log2 Fold Change respectively. We then used the “MSigDB_Hallmark_2020” gene sets defined by the Molecular Signatures Database (MSigDB)^63^, representing 50 well-defined biological processes. We applied enrichr on responders and non-responders separately and considered a significance cutoff of 0.05 for the adjusted p-values (Table S2). This analysis was conducted on the integrated datasets of both tumor and blood samples separately (Supplementary Figures S4-5).

### Detection of a robust expansion response signature

In order to identify a response signature of expanded CD8^+^ T cells that is robust and predictive of response across datasets, we utilized 9 discovery datasets having samples of both responders and non-responders^2,3,5,8,14,15,18,19^. We then conducted differential gene expression between expanded CD8^+^ T cells from both groups in each dataset separately. We used Scanpy’s *‘scanpy.tl.rank_genes_groups’* function with *‘method = wilcoxon’* and considered significant genes as those that have a non-zero expression in at least 20% of the cells (for at least one of the tested groups - responders/non-responders), with an adjusted p-value < 0.05 and log2 Fold Change > 0.5 for responders (or log2 Fold Change < -0.5 for non-responders). We further considered genes such that the difference of the non-zero expression was larger than 10% of the cells between responders and non-responders. The full lists of differentially expressed genes per dataset are summarized in Table S3.

For each gene, we counted the number of datasets (out of the 9 discovery datasets), where it was considered as significant in expanded CD8^+^ T cells for responders/non-responders. The genes were then ordered according to the number of datasets where they were considered as significant, as well as according to their rank calculated as: −log_10_(𝑀𝑒𝑎𝑛_𝑝_𝑣𝑎𝑙𝑢𝑒_𝑎𝑑𝑗_) ·𝑀𝑒𝑎𝑛_log_2_(𝐹𝑜𝑙𝑑𝐶ℎ𝑎𝑛𝑔𝑒), such that the mean was applied for each gene using the datasets where it was considered as significant in responders and non-responders separately. This ordering defined a genetic signature of expansion markers in responders and non-responders and is summarized in Table S3.

We then used these two signatures and scored each expanded CD8^+^ T cell based on the number of expressed genes (log_2_(𝑛𝑜𝑟𝑚𝑎𝑙𝑖𝑧𝑒𝑑_𝑐𝑜𝑢𝑛𝑡𝑠 + 1) > 1) out of the top K response markers of each signature, yielding two scores for each expanded T cell. Then, each T cell was classified as ‘favorable’ or ‘unfavorable’ based on the majority vote of the two scores. In cases where both scores were tied in a single cell, it was classified as ‘favorable’.

Finally, we computed a score per sample by taking the ratio between the number of cells classified as ‘favorable’ out of the total number of cells classified as ‘favorable’ or ‘unfavorable’, yielding a score per sample between 0 and 1. T cells demonstrating an expression of zero markers from both lists of markers were excluded from this analysis. We performed this process for different values of K, ranging 2,…,20. For each K, we calculated the Receiver Operating Curve (ROC) and the Area Under the Curve (AUC) score, as well as the p-value. Best results were achieved for K = 5, 6 (Figure 2B, Table S3). Further reviewing this solution, we chose K = 6 due to better performance in individual datasets. A two-sided Wilcoxon rank sum test was used to calculate the p-value, comparing the ratio score between responders and non-responders per dataset (Figure 2, Supplementary Figure S6). P-values were corrected for multiple hypothesis using the Benjamini-Hochberg false discovery rate^61^. It is important to note that samples that did not contain expanded CD8^+^ T cells that potentially target cancerous neoantigens (as described above) were not used for this analysis.

### Validation of the expansion response signature on additional paired scRNA/TCRseq datasets

We first validated our devised response signature using two cohorts of ICI treated breast cancer patients by Bassez et al.^16^, considering only samples containing expanded CD8^+^ T cells (Figure 2J). The patients in these datasets lack response status, though they were originally annotated for treatment-related expansion at the level of the single patient. These datasets were included in our study in order to refine the single-cell clusters, cNMF programs and the clonal gene trajectory described below. Besides these datasets, we used additional dataset of NSCLC patients containing only non-responders^17^ (Supplementary Figure S6D), and two additional external datasets with paired scRNA/TCRseq of cancer patients having response status (Table S1). The first includes De-novo glioblastoma patients treated with ICI and CAR therapy, all failed to respond^40^. The second consists of melanoma patients treated with adoptive cell therapy, having both responders and non-responders^41^. Both scRNAseq and scTCRseq from these two datasets were preprocessed as described above. We then focused on expanded CD8^+^ clones as described above and tested the significance of our devised response signature between responders and non-responders, using a two-sided Wilcoxon rank-sum test (Figure 2K, Supplementary Figure S6D). Only samples containing expanded CD8^+^ T cells were considered.

### Validation of the expansion response signature on sorted bulk samples

As our signature is T-cell related, we made additional validation on bulk samples of treatment-naive NSCLC patients sorted for T cells^42^ (Table S1). We used the original labels regarding CD8^+^ T cell infiltration per sample as provided by the authors (TIL^hi^, TIL^int^ or TIL^lo^), and processed the count matrix using the default pipeline of the DESeq2 package^64^. For each gene (out of the 12 favorable and unfavorable genes), we calculated the median expression in normalized counts across all samples, and scored each sample by the amount of favorable/unfavorable genes that were expressed above the median. This process resulted with a ratio per sample, indicating how many favorable genes were highly expressed in that sample out of all the favorable and unfavorable genes that were highly expressed. We then applied a two-sided Wilcoxon rank-sum test between these ratios in samples classified as TIL^lo^ and the rest of the samples (Figure 2L, Supplementary Figure S6E).

### Identifying cNMF programs and their clonal changes following administration of therapy

We applied cNMF^43^ on the combined raw count matrix of all the single-cells passing QC from both tumor and blood samples, using all of the 12,407 genes that passed our QC and existed across all datasets. We used 200 NMF replicates for each K and tested the results for K = 2,…,30. Considering the error and stability values for each K, we selected K = 19 as the optimal solution. Further reviewing this solution, we filtered out seven programs that were active (considering the max usage value) in less than 1% of the cells, leaving us with 12 programs overall (Figure 3A, Supplementary Figure S8A, Table S4). Each program was annotated based on its top-ranked genes, and four programs having non-distinct annotations underwent an over-representation analysis as described above, using their top 100 genes (Table S4). To identify significant clonal changes of programs following administration of therapy, we focused only on expanded CD8^+^ clones that do not target a known non-cancerous antigen as described above.

We then analyzed clones that were shared between pre and post samples of the same patient that were also expanded in both time-points (termed ‘persistent’ clones). We averaged the usage of each program per clone to obtain the mean usage value in every clone for both time-points. In cases where a single patient had multiple samples taken following therapy containing the same expanded clone, we averaged the program usage across all the cells from the same clone in those samples. In addition, two NSCLC patients from Liu et al.^3^ had multiple biopsies taken following therapy with different response statuses. In these cases, we considered the pre/post pairs that were taken from the same biopsy site and had the same response status. Overall, 33 responders and 17 non-responders had persistent expanded CD8^+^ clones existing both in their pre and post samples. For each patient, we focused on the top 5 expanded clones that existed in both pre/post samples based on the mean clone size of each clone across all samples of the same patient, where the clone was expanded. In cases where less than 5 expanded clones were shared between the two time-points, we considered the clones that did exist. In cases where the fifth top largest clone had the same mean clone size as the next clones in order (i.e. the 6^th^ clone etc.), we included these clones as well using the function *‘pandas.DataFrame.nlargest’* with the argument *‘keep = all’*. We then conducted paired two-sided Wilcoxon signed-rank test between the mean usage of each program for these clones per patient in the two treatment time-points. We conducted this test for responders and non-responders separately, and corrected the obtained p-values for multiple hypothesis using the Benjamini-Hochberg false discovery rate^61^. This process was conducted for tumor and blood samples independently (Figure 3, Supplementary Figures S7-8), and was also tested for shared clones that appeared in both blood and tumor biopsies of the same patient (Supplementary Figure S13). For analysis of shared clones appearing both in tumor and blood, we used the average values in tumor/blood samples in cases where the same clone was expanded in more than one tumor/blood sample of the same patient.

We repeated this analysis for clones that were expanded in tumor samples following therapy (termed “de-novo clones”), across all patients having longitudinal biopsies (Supplementary Figure S12). We focused on the top 5 expanded clones per sample using the function *‘pandas.DataFrame.nlargest’* with the argument *‘keep = all’* and extracted only the de-novo clones out of the top expanded clones. It is important to note that for this analysis we used only samples having de-novo clones among their top 5 expanded clones. We then conducted a two-sided Wilcoxon rank-sum test between the mean usage of each program per clone between de-novo clones in responders and non-responders. P-values were then corrected for multiple hypothesis using the Benjamini-Hochberg false discovery rate^61^ (Supplementary Figure S12).

### Defining the set of metabolic genes used to identify metabolic cNMF programs

In order to further focus on metabolic genes independently, we used a set of metabolic genes and pathways that is based on the Recon 2 metabolic reconstruction^45^. We used the set of 1,193 metabolic genes from 96 metabolic pathways that exist in our integrated dataset (Table S4). The glycolysis pathway as defined in Recon 2 is composed of 53 genes (that passed our QC) and includes also gluconeogenesis-related genes. Therefore, to specifically study glycolysis in our analysis, we separately defined a “Glycolysis” pathway which includes only 19 central genes (Table S4), as was done similarly in our previous work^35^.

We then similarly applied cNMF^43^ as described above, using all of the 1,193 metabolic genes^45^ that passed our QC and existed across all datasets. Considering the error and stability values for each K, we selected K = 8 as the optimal solution. Further reviewing this solution, we filtered out two programs that were active (considering the max usage value) in less than 1% of the cells, leaving us with 6 programs overall (Supplementary Figure S9, Table S4). A paired two-sided Wilcoxon signed-rank test was then conducted between the mean usage of each program for the top 5 expanded persistent clones in the two treatment time-points as described above, and for clones that were shared between tumor and blood samples (Supplementary Figures S9, S14). P-values were corrected for multiple hypothesis using the Benjamini-Hochberg false discovery rate^61^.

### Clonal pseudobulk analysis

To study the clonal pseudo-temporal dynamics, we analyzed each single expanded CD8^+^ clone in a pseudobulk manner based on the mean expression of the genes across all the single cells from the same clone per sample. We again focused on expanded CD8^+^ clones that potentially target cancerous neoantigens as described above, and focused on 7,945 expanded clones that originated in tumor samples. For data visualization, we applied the same preprocessing steps for the pseudobulk expression matrix as described above, and removed batch effects using Batch Balanced K-Nearset Neighbors (BBKNN)^59^ with a *‘batch_key’* of *‘study’* in order to remove batch effects between clones from different single-cell studies (Figure 4A).

### Clonal gene trajectory

In order to identify pseudo-temporal dynamics of single clones, we used GeneTrajectory^49^ – an approach that identifies trajectories of genes rather than trajectories of cells. This method outperformed cell-trajectory methods in recovering the gene order for both cyclic and linear processes^49^. We used genes having a non-zero expression in at least 5% of the clones and at most in 90% of the clones, out of the top 1000 highly variable genes across these clones.

We used the *‘gene_trajectory.get_graph_distance’* function with *‘k=20’* of kNN and followed the standard protocol provided by the authors, achieving a single gene-trajectory for the analyzed expanded CD8^+^ clones (Table S5). The gene trajectory was then projected over the UMAP of the pseudobulk clones using the *‘add_gene_bin_score’* function with *‘n_bins’* = 1 (Figure 4A). We then used the function *‘plot_gene_trajectory_2d’* in order to generate a two-dimensional representation of the genes with their sequential manner (Figure 4B).

### Clonal exhaustion, memory and metabolic scores

We used Scanpy’s function *‘scanpy.tl.score_genes’* with its default parameters to determine the following scores per clone, as was similarly done previously in the single-cell level by Oliveira et al.^39^: memory score (*IL7R*, *SELL*, *CCR7*, *CD28*, and *TCF7*) and exhaustion score (*PDCD1*, *HAVCR2*, *TIGIT*, *CTLA4*, *LAG3*, and *TOX*).

To compute a metabolic pathway score for each pathway in every single clone, we applied similar steps to those done in our previous work^35^: we first calculated for every single cell from each clone the amount of expressed genes related to each Recon 2 metabolic pathway^45^ (Table S4), out of the total number of genes in that pathway. This process resulted with a continuous score between 0 and 1 for each pathway in each single cell. Specifically, for each metabolic pathway K, the metabolic pathway score of each single cell was calculated such that:

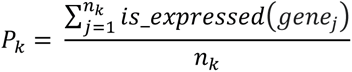

Where 𝑖𝑠_𝑒𝑥𝑝𝑟𝑒𝑠𝑠𝑒𝑑: 𝑥 → {0, 1} and defined as:

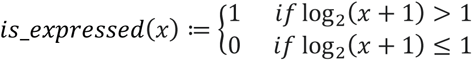

𝑥 is the expression level in normalized counts, 𝑛_𝑘_is the total number of genes in metabolic pathway K, and the sum is applied to all of the genes such that 𝑔𝑒𝑛𝑒_𝑗_ ∈ 𝑝𝑎𝑡ℎ𝑤𝑎𝑦_𝑘_

We then averaged the metabolic pathway scores for all the single cells in every clone per sample, to achieve a single score per clone in every sample, for every metabolic pathway.

We then tested the Spearman correlation of each metabolic score with the gene trajectory (Table S5). P-values were corrected using the Benjamini-Hochberg false discovery rate^61^. When focusing on metabolic pathways with a median clonal score larger than 0.2, ‘Glycolysis’ and ‘Oxidative phosphorylation’ were the two top pathways correlated with the gene trajectory (Figure 4A, Table S5). The Spearman correlation was also tested between the clone size and the gene trajectory (Figure 4C).

### Differences in gene trajectory of dying, de-novo or persistent CD8^+^ clones

Using CD8^+^ expanded clones from patients having longitudinal biopsies, we classified each CD8^+^ clone as dying/de-novo/persistent as described above. We focused on the top 5 expanded CD8^+^ clones per sample as described above and used a two-sided Wilcoxon rank-sum test in order to test the differences in gene trajectory between dying and de-novo clones, as well as persistent clones before and following treatment in responders and non-responders (Figure 4D).

### Analysis of intra-tumoral CD8^+^ clones shared with blood samples

In order to study differences between intra-tumoral clones that are shared between matched blood and tumor samples vs. those that are restricted to the tumor, we used three datasets containing patients having both tumor and blood samples^5,15,19^ (Figure 5A, Table S1). We focused on intra-tumoral expanded CD8^+^ clones that potentially target cancerous neoantigens as described above, and labeled them as “shared” if they were also expanded in blood samples, or as “not-shared” if they were expanded solely in the tumor, regardless of treatment time-point. We then tested for differences in the activity of cNMF programs between both groups, using the mean usage of each program per clone. We applied a two-sided Wilcoxon rank-sum test and corrected the obtained p-values using the Benjamini-Hochberg false discovery rate^61^ (Figure 5B, Supplementary Figure S15A). The abundance of intra-tumoral shared clones was then calculated per sample as the number of shared clones out of all the expanded CD8^+^ clones, and was tested between responders and non-responders using a two-sided Wilcoxon rank-sum test (Figure 5D). Considering the mean usage of the cNMF programs across all the single cells per clone, we calculated the fraction of clones having max usage of each program in both shared and non-shared clones (Figure 5E).

In order to generalize our findings to additional datasets having tumor samples without matched blood samples, we used the max-usage of the FGFBP2-NKG7 program as a potential marker of intra-tumoral shared clones (due to its high abundance in shared clones). We considered additional seven datasets^2,3,8,14,17,18^ containing only tumor samples and classified each expanded clone based on its max usage of the cNMF programs. We then calculated per sample the fraction of expanded CD8^+^ clones having max usage of the FGFBP2-NKG7 program, out of all the expanded CD8^+^ clones. We conducted a two-sided Wilcoxon rank-sum test between these ratios in responders and non-responders (Figure 5F).

### Validation of the max FGFBP2-NKG7 clonal ratio on additional paired scRNA/TCRseq dataset

We utilized additional dataset of MCC patients treated with ICI^50^. Both scRNAseq and scTCRseq were preprocessed as described above, resulting with 31 samples from 27 patients passing QC. One sample was removed for having only one cell passing QC, leaving us with 30 samples from 26 patients overall (Supplementary Figure S16A-B). We applied NMFproj^65^ in order to reflect our cNMF programs upon this new dataset, following the default pipeline suggested by the authors (Supplementary Figure S16C). We then focused on expanded CD8^+^ clones with high potential to target cancerous neoantigens as described above and calculated per sample the fraction of expanded clones having max usage of the projected FGFBP2-NKG7 program, out of all the expanded clones. We conducted a two-sided Wilcoxon rank-sum test between these ratios in responders and non-responders (Figure 5G).

## Supplemental information titles and legends

**Document S1. Figures S1-S16.**

**Table S1. Summary of utilized paired scRNA/TCRseq datasets,** Related to Figure 1.

**Table S2. List of differentially expressed genes and pathway enrichment analysis,** Related to Figures 1, S1-5.

**Table S3. Analysis of study-wise response signature of expanded T cells,** Related to Figures 2, S6.

**Table S4. cNMF programs and metabolic genes,** Related to Figures 3-5, S7-16.

**Table S5. Gene trajectory analysis of CD8^+^ expanded clones,** Related to Figures 4, S11.

